# Datanator: an integrated database of molecular data for quantitatively modeling cellular behavior

**DOI:** 10.1101/2020.08.06.240051

**Authors:** Yosef D. Roth, Zhouyang Lian, Saahith Pochiraju, Bilal Shaikh, Jonathan R. Karr

## Abstract

Integrative research about multiple biochemical subsystems has significant potential to help advance biology, bioengineering, and medicine. However, it is difficult to obtain the diverse data needed for integrative research. To facilitate biochemical research, we developed Datanator (https://datanator.info), an integrated database and set of tools for finding *clouds* of multiple types of molecular data about specific molecules and reactions in specific organisms and environments, as well as data about chemically-similar molecules and reactions in phylogenetically-similar organisms in similar environments. Currently, Datanator includes metabolite concentrations, RNA modifications and half-lives, protein abundances and modifications, and reaction rate constants about a broad range of organisms. Going forward, we aim to launch a community initiative to curate additional data. Datanator also provides tools for filtering, visualizing, and exporting these data clouds. We believe that Datanator can facilitate a wide range of research from integrative mechanistic models, such as whole-cell models, to comparative data-driven analyses of multiple organisms.

## 1. Introduction

Integrative research about multiple biochemical subsystems has significant potential to help advance biology, bioengineering, and medicine. For example, whole-cell models that account for all of the biochemistry in cells could help scientists conduct experiments in silico, help physicians personalize medicine, and help engineers design microbes.^1, 2^

Historically, it has been challenging to obtain the diverse and extensive data needed to analyze multiple subsystems. For example, Resource Balance Analysis (RBA) requires data about the composition of cells and macromolecular complexes, the abundances of enzymes, and the maximum rates of reactions.^3^ Whole-cell models require even more data, such as the concentration of each metabolite, the abundance and turnover of each RNA and protein, and the rate law of each reaction.

New experimental technologies have begun to produce the data needed for integrative research. For example, RNA-seq has enabled transcriptome-wide profiles of RNA abundances and half-lives, tandem mass spectrometry has enabled proteome-scale profiles of protein abundances and half-lives, and liquid chromatography-mass spectrometry has enabled metabolome-scale profiles of metabolite concentrations. Furthermore, this data is increasingly accessible. For example, ArrayExpress contains over 400,000 RNA expression profiles.

Several investigators have begun to use this data to conduct more integrative analyses of biochemical networks. For example, Sanchez et al.^4^ used proteomic data and reaction rate parameters to develop a constraint-based model of the gene expression and metabolism of yeast; Thiele et al.^5^ used genetic information and other data to build a constraint-based model of the transcription, RNA modification, translation, complexation, and metabolism of *Escherichia coli*; Goelzer et al. used metabolomic and proteomic data to develop an RBA model of 72 subsystems of *Bacillus subtilis*;^3^ and we and others used a wide range of data^6^ to develop a hybrid model of 28 subsystems of *Mycoplasma genitalium*.^7^ However, even the most extensive studies have only used a fraction of the available data. Encouraged by these successes, we believe that more comprehensive analysis is possible with more data.

Although extensive data is available, a recent community survey revealed that obtaining data is one of the biggest bottlenecks to integrative research.^8^ One fundamental barrier is our limited ability to experimentally characterize biochemistry. For example, the most comprehensive metabolomics datasets only capture a small fraction of the metabolome. Long-term, the community must develop new measurement technologies.

It is also difficult to utilize the existing data: the existing data is siloed in different databases and articles for different types of data and organisms; the existing databases and articles use different formats, identifiers, and units (e.g., PAXdb provides data in TSV format, whereas SABIO-RK provides data in SBML format); the APIs to the existing databases have different interfaces; and the existing databases provide limited tools for finding data for similar biology for imputing missing information. This heterogeneity makes it difficult to compare and integrate data from multiple sources.

We believe these secondary challenges can be addressed with practical computational solutions. To address these challenges, we developed Datanator (https://datanator.info), an integrated database of molecular data and a single web interface and API for finding a broad range of quantitative and categorical data about a wide range of organisms.

To start, we have aggregated several key types of data, including metabolite concentrations, RNA modifications and half-lives, protein abundances and modifications, and reaction rate parameters from several databases and articles. Going forward, we aim to launch a community initiative to curate additional data. To help researchers search, compare, and integrate this heterogeneous data, we have normalized the annotation, units, and representation of the data.

On top of this database, Datanator provides tools for finding relevant data for research. Like other databases, Datanator provides tools for finding data about specific molecules and reactions in particular organisms and environments. In the absence of direct data, Datanator also provides unique tools for building *clouds* of data that include chemically-similar molecules and reactions in phylogenetically-similar organisms and similar environments. For example, Datanator can find data about the concentration of ATP and similar metabolites in *E. coli* or other enterobacteria in M9 or other minimal media. These clouds can help researchers impute missing information. Datanator also provides tools for filtering these clouds, charting their distribution, and exporting them to files.

We believe that Datanator can facilitate a wide range of research. For example, Datanator can help investigators find information for comparative analyses of multiple organisms and tissues, multi-dimensional analyses of relationships within biochemical networks, and integrative mechanistic models of multiple subsystems such as whole-cell models.

Here, we describe the content of Datanator and the search and visualization tools. In addition, we summarize how we implemented Datanator, compare Datanator to several existing databases, outline the types of research that Datanator can facilitate, and discuss how we plan to enhance Datanator.

## 2. Integrated database of molecular data

Because research often requires many types of data, Datanator includes several types of quantitative and categorical data. Currently, this includes 3,841 measurements of the concentrations of 1,621 metabolites aggregated from ECMDB,^9^ YMDB,^10^ and several articles; 589 reconstructed RNA modification profiles aggregated from MODOMICS;^11^ 75,836 measured RNA half-lives aggregated from several articles; 2,634,941 measurements of the abundance of 846,970 proteins aggregated from PaxDB;^12^ 4,083 reconstructed protein modification profiles aggregated from PRO;^13^ and 61,734 measurements of the rate parameters of 37,858 reactions aggregated from SABIO-RK^14^ (Figure 1a). In addition, we are working to include nearly 1,000,000 measurements of RNA localizations from RNALocate,^15^ lncATLAS,^16^ and one article; over 100,000 measured and predicted protein localizations from eSLDB,^17^ the Human Protein Atlas,^18^ PSORTdb,^19^ and Sub-Cell;^20^ more than 50,000 measurements of protein half-lives from nine publications; more than 200,000 additional measurements of reaction parameters from BRENDA;^21^ and reaction flux measurements for 36 organisms from CeCaFDB.^22^

**Figure 1.**
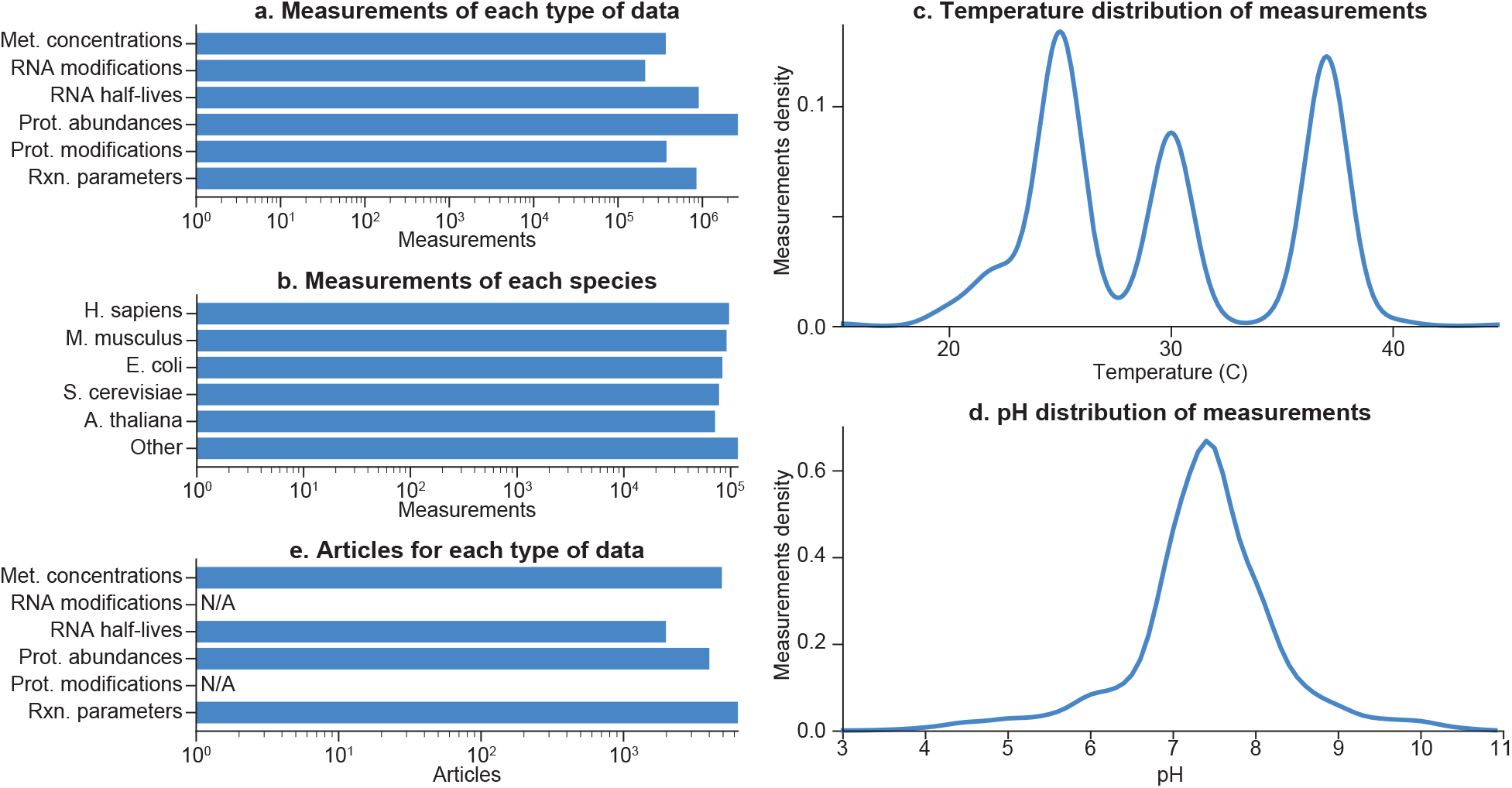
Datanator encompasses several key types of molecular data about a wide range of organisms in a broad range of environments. **(a)** Number of measurements of each type of data. **(b–d)** Distributions of the genotype and environment of the measurements in the database. **(e)** Number of articles that Datanator integrates data from for each type of data.

In total, Datanator currently includes data for 1,030 organisms (Figure 1b) across a wide range of environmental conditions (Figure 1c,d) from over 8,000 articles (Figure 1e).

## 3. Tools for searching the sea of data

Datanator also provides unique tools for searching for relevant data for research. Like other databases, Datanator provides basic functionality for finding data about specific molecules and reactions in particular organisms and environments. On top of this, Datanator provides tools for assembling *clouds* of related data – ensembles that also encompass measurements of similar molecules and reactions in similar organisms and environments. In the absence of direct measurements, these clouds can help researchers impute missing information.

### Basic searching for specific measurements

Users can use the search form (Figure 2a) to query for data about a specific metabolite, gene (RNA or protein), or reaction in a specific organism. Users can search for data about molecules and reactions by their names, identifiers (e.g., ChEBI id), chemical structures (e.g., InChI representation), and descriptions. Datanator recognizes all organisms in the NCBI Taxonomy database. After submitting the form, Datanator presents three lists of search results for metabolites, genes, and reactions, each ranked by their relevance to the query (Figure 2b–d). Users can click on each result to obtain tables of measured properties (e.g., concentration, half-life) of the selected molecule or reaction (Figure 2e).

**Figure 2.**
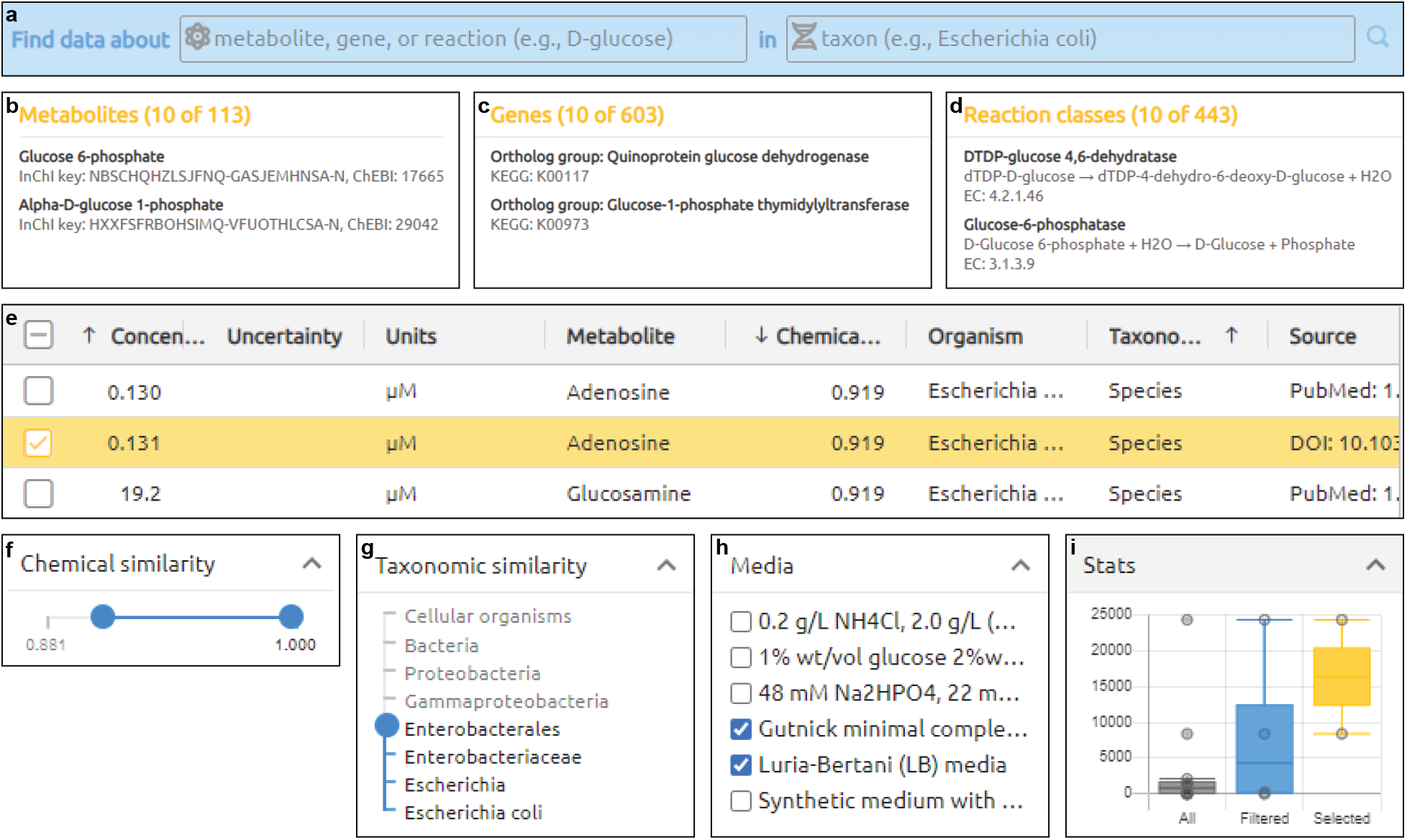
Datanator provides interactive tools for creating and visualizing clouds of data centered on specific molecules and reactions in specific organisms and environments. (**a**) Search form. (**b–d**) Search results grouped by class (metabolites, genes, or reaction). (**e**) Tables of clouds of potentially relevant measurements about a molecule or reaction. (**f–h**) Tools for filtering the data clouds. (**i**) Charts for visualizing the distributions of the data clouds.

### Advanced searching for clouds of similar measurements

While the amount of data has exploded over the last decade, we still do not have data about every molecule and reaction in every organism and environment. Instead, researchers often must impute missing information from data about similar molecules and reactions in related organisms and environments. However, it is difficult to find data about similar biology. For example, searching Google for data about ATP synthase or its orthologs in *E. coli* or other Enterobacteria such as *Klebsiellas* requires combinatorially-many searches for each ortholog in each Enterobacterial species.

To help researchers conduct research even without direct measurements, Datanator provides tools for assembling *clouds* of data around a specific molecule or reaction in a particular organism and environment. For metabolites, Datanator identifies measurements of chemically-similar metabolites (Tanimoto coefficient ≥ 0.65). For genes, Datanator identifies measurements of the same KEGG Orthology group. For reactions, Datanator identifies measurements of the same Enzyme Commission (EC) class. Datanator identifies measurements from all organisms and environments.

Datanator displays these clouds using the data tables described above (Figure 2e). To help users understand these clouds, the data tables include columns that describe the similarity of each measurement to the search query. For metabolites, Datanator displays the Tanimoto coefficient between the target and measured metabolites. Datanator indicates the phylogenetic relevance of each measurement by displaying the taxonomic rank of the most recent canonical-ranked common ancestor (e.g., species, genus) between the target and measured organisms.

To help users narrow these data clouds to the most relevant measurements, Datanator provides sliders and select menus for filtering measurements by their metadata (Figure 2f–h). This includes a slider for filtering metabolite concentration measurements by their Tanimoto coefficient, a slider for filtering measurements by their taxonomic distance, sliders for filtering measurements by their temperature and pH, and a select menu for filtering measurements by their growth media.

In addition, Datanator provides users checkboxes for creating data clouds manually (Figure 2e). These fine-grained controls can help users create clouds that only include the data that they believe are relevant to their research.

Finally, Datanator provides buttons for exporting data clouds to CSV and JSON files.

### Programmatic searching via a REST API

Users can also programmatically query Datanator via a REST API. The API makes it easy for users to obtain data about multiple molecules or reactions in multiple organisms.

## 4. Tools for visualizing data clouds

To help researchers interpret their data clouds, Datanator provides charts of the distributions of all of the potentially relevant data, the filtered measurements, and the selected measurements (Figure 2i). Users can mouse over each measurement to highlight it in the corresponding data table.

## 5. Implementation

Datanator consists of a database of molecular data, an OpenAPI-compliant REST API for querying the database, and a web application for graphically exploring the database.

### Database construction

We assembled the Datanator database in several steps. First, we used Google, Google Scholar, and Microsoft Academic to find potential sources for each type of data. Second, for each kind of data, we decided to focus on the largest databases. In the absence of an existing database, we decided to focus on articles that report genome-scale measurements.

Third, we downloaded and parsed each source. Where available, we used file downloads and REST APIs. For example, we downloaded PaxDB as an archive of TSV files, and we downloaded the Protein Ontology as an OBO file. For databases that do not provide a download or complete API, we wrote scripts to scrape and parse their websites.

Fourth, we normalized each measurement. We normalized the subject of each measurement (molecule or reaction) by mapping it to one of several namespaces. We used Open Babel^23^ to map each metabolite to the InChI representation of its structure, mapped each mRNA and protein to its UniProt id, and mapped each reaction to the InChI representations of its participants. We used BpForms^24^ to normalize the representation of each RNA and protein modification profile. We normalized the taxonomic context of each measurement by using the ETE toolkit^25^ to map each organism to its NCBI Taxonomy id. We used the pint Python package to normalize the units of each measurement. Lastly, we combined the measurements into a single MongoDB database.

To enable us to pull updates to each source into Datanator going forward, we implemented the database construction as a repeatable script.

### REST API

We implemented the API using Python, connexion, pymongo, and Elasticsearch.

### Web application

We implemented the web application using the React framework and several packages, including ag-Grid (data tables), Blueprint (search form), and ChartJS (charts).

### Testing

We tested the database construction and REST API using unittest. We tested the web application using Cypress and Jest.

## 6. Comparison to existing databases

Numerous resources help researchers find data, including databases for individual types of data and model organism, genome, pathway, and federated databases. While several of these resources overlap with Datanator, Datanator uniquely contains multiple types of quantitative and categorical molecular data for multiple organisms. When direct measurements are unavailable, Datanator also provides unique tools for finding data about similar biology.

### Data type-specific databases

Although most of Datanator’s data is available from other databases such as ECMDB and PaxDB, it is challenging to use these databases together because they provide data via different interfaces in different formats, and they annotate their data with different identifiers and ontologies. In addition, these databases provide few tools for obtaining data about similar biology for imputing missing information. Datanator makes their data more accessible by providing a central, consistent portal to the data and a complete API. Datanator also provides tools for building clouds of their data. Additionally, Datanator has more metabolite concentrations than ECMDB or YMDB because it combines ECMDB, YMDB, and several articles.

### Model organism databases

Several model organism databases have overlapping content with Datanator. EcoCyc contains reaction kinetic parameters for *E. coli*.^26^ SubtiWiki contains RNA and protein expression data for *B. subtilis*.^27^ However, these databases focus on genetic and relational data, and they each only contain data for one organism. The CCDB (*E. coli*, 28), MyMPN (*Mycoplasma pneumoniae*, 29), and WholeCellKB (*M. genitalium*, 6) contain multiple types of data. Datanator improves upon these databases by incorporating data for multiple organisms and providing more powerful search tools.

### Genome and pathway databases

Several genome and pathway databases also have overlapping content with Datanator. For example, Reactome contains RNA and protein expression data^30^ and UniProt contains data about protein modifications.^31^ However, these databases focus on genetic and relational information, such as the locations of promoters and their regulators. These databases have limited quantitative information, such as concentrations.

### Federated databases

Like Datanator, federated databases such as BioMart^32^ and Omics DI^33^ are also central portals for discovering multiple types of data. While both databases contain more data than Datanator, their content has little overlap with Datanator. Furthermore, the content of Datanator is more integrated than that of BioMart and Omics DI. This integration makes it easier for researchers to search Datanator and use its data.

## 7. Use cases

Through helping researchers obtain data, we believe that Datanator can assist a wide range of tasks such as comparative analyses of multiple tissues and organisms, multi-dimensional analyses of molecular networks, calibrating mechanistic models, and validating conclusions with independent data.

### Multi-dimensional analyses of biochemical networks

Because Datanator contains multiple types of data, Datanator is a good source of data for analyzing relationships among molecules, reactions, and their chemical properties. For example, Datanator could provide data for analyzing the correlation between the abundances of enzymes and their maximum reaction velocities, classifying reactions as bottlenecks by comparing metabolite concentrations and enzyme-metabolite affinities, or learning a regression model of the half-lives of RNAs from their sequences and modifications.

### Calibrating mechanistic models

For similar reasons, Datanator is also a good source of data for calibrating mechanistic models. For example, Datanator could provide protein abundances to constrain missing initial conditions; provide metabolite concentrations to construct a more comprehensive biomass equation for a flux balance analysis (FBA) model; or provide organism, tissue, or cell-type-specific protein abundances for recalibrating a model to create variants for different organisms or cell types. In particular, Datanator is ideal for integrative modeling of multiple biochemical subsystems such as whole-cell models. For example, Datanator could provide protein abundances and maximum reaction velocities to expand an FBA model to capture gene expression and further constrain the model.^4^

### Validating computationally-generated conclusions

In the absence of an experimental collaboration, computational scientists often validate their conclusions with independent data from articles. However, this is often time-consuming because this requires finding data about specific molecules or reactions in particular organisms. By making it easier to find data about specific biology, Datanator empowers computational scientists to validate their own conclusions.

## 8. Discussion

In summary, Datanator is a unique portal for obtaining several key types of quantitative and categorical molecular data. The foundation of Datanator is an integrated database of several types of data about numerous organisms. This includes the first collection of multiple studies of RNA half-lives. Datanator is also the first database to contain the concentrations of metabolites in multiple organisms. To help researchers use this data, Datanator provides unique tools for assembling clouds of data centered on specific molecules and reactions in particular genotypes and environments. In addition, Datanator provides charts for visualizing the distribution of these clouds. These clouds can help users obtain informative data even in the absence of direct measurements. We anticipate that Datanator will help researchers build integrative models of multiple biochemical subsystems, including models of entire cells. We believe that Datanator is also a valuable source of data for multi-dimensional analyses of biochemical networks and comparative studies of multiple organisms.

### Community initiative to curate additional data

Going forward, we aim to launch a community initiative to curate additional data. To facilitate community contributions, we aim to make it easy to submit data. To start, we have developed a single format for representing any type of data and a tool for validating data sets. This format outlines how investigators can submit data. Already, this is enabling several high school students to contribute data about protein localizations and half-lives. Ultimately, we plan to accept data through a simple form. We welcome suggestions via GitHub issues and pull requests or by joining our team.

### Additional search and analysis tools

To further help users use the Datanator data, we plan to develop additional search and analysis tools. For example, we aim to create charts of the global distribution of each type of data; implement a tool for extracting multi-dimensional data clouds such as clouds of the expression and half-lives of mRNA and their proteins; and develop finer-grained filters for data about similar RNAs, proteins, and reactions based on the sequence similarity of RNAs and proteins and the structural similarity of the participants of reactions.

### Higher-level tools for integrative modeling

In addition, we aim to develop higher-level tools for leveraging Datanator for integrative modeling. For example, we aim to build tools that further automate the process of using data from Datanator to constrain the values of missing model parameters, recalibrate models to represent different cell types, and further constrain FBA models with enzyme abundances and reaction parameters.

## 9. Availability

The application and API are available at https://datanator.info and https://api.datanator.info along with a tutorial and documentation. The data and software are openly available under the CC BY-NC-ND and MIT licenses, respectively.

## 10. Acknowledgements

We thank Yin Hoon Chew, Paul Lang, Wolfram Liebermeister, and the Center for Reproducible Biomedical Modeling for feedback.

